# WillCO_2_st: leveraging freely available real-time carbon forecasting for research emissions reporting

**DOI:** 10.64898/2026.01.30.702771

**Authors:** William V. Smith, Stefan R. Pulver

## Abstract

An increasing number of research institutions and funding bodies are making sustainability a focal point. The rapid rollout of auditing processes, such as the Laboratory Efficiency Assessment Framework (LEAF), and changing institutional policies, are creating demand for sustainability accounting upon researchers. Despite the increasing expectations on researchers to actively report their sustainability and demonstrate improvement, easy-to-use, and accurate reporting tools for electricity-associated emissions (scope 2) accounting are not widely available. Here, we created WillCO_2_st - a free program that enables fast, publication-read sustainability reports. WillCO_2_st makes use of real-time, region-specific conversion factor data freely available through resources like the Carbon Intensity and Electricity Maps API to create experimental carbon footprinting estimates using metadata from saved experimental files. Data can be simply click-and-dragged into WillCO_2_st to create an experimental sustainability audit within minutes. Further, we exemplified the utility of WillCO_2_st’s live-nudging feature by quantifying the retrospective theoretical reduction in electricity-driven emissions from shifting experimental timing to times of day with lower carbon intensities. Overall, WillCO_2_st eases time and resource burden on researchers and serves as a model for how researchers can effectively and accurately engage with research sustainability auditing.

**Highlights:**

⍰ WillCO_2_st is a free software platform that enables fast, publication-ready carbon emissions reporting for papers, grants, and internal audits.
⍰ WillCO_2_st leverages various energy-to-emissions database to provide real-time, region-specific estimations of carbon intensity for the upcoming 48 hours, nudging users to switch to lower carbon times for research activities.
⍰ WillCO_2_st integrates CodeCarbon to automatically audit Python-based coding scripts for experimental and computational-wide sustainability audits.
⍰ Partial conformity (21%) to switching experimental time would have disproportionately produced a large carbon saving (50%) in our most recent publication’s experimental carbon footprint.

## Introduction

Institutions and research funding bodies are becoming increasingly climate-conscious, and this places demands on researchers to plan, foster, and demonstrate sustainable research practices. A larger number of UK universities (e.g., Russell Group) and research institutions (e.g., UK Research & Innovation (UKRI), Wellcome Trust) have signed the Concordat for the Environmental Sustainability of Research and Innovation Practices [1], creating explicit commitments to progressively embed awareness of environmental sustainability across research and innovation communities, and foster direct action to mitigate the emissions from research. Currently, many UK universities use auditing frameworks like the Laboratory Efficiency Assessment Framework (LEAF) to evaluate, mitigate, and signal research sustainability [2], [3]. The drive for research sustainability is being propelled by major funding bodies that are pushing researchers to be conscious and proactive in sustainability innovation and reporting. Major public funders across the world like UKRI [1], Wellcome Trust [4], Cancer Research UK (CRUK) [5], National Institute for Health and Care Research (NIHR) [6], British Heart Foundation (BHF) [7], the Royal Academy of Engineering [8], Horizon Europe [9], Swiss National Science Foundation [10], Deutsche Forschungsgemeinschaft [11], National Science Foundation [12], National Institute of Environmental Health Sciences [13], Australian Research Council [14], Research Ireland [15], Marie Sklodwska-Curie Actions and Green Charter [16], and Royal Society of Chemistry (RSC) [17] have explicitly made sustainability a focal element in funding themes, with some bodies requiring action on sustainability in order to receive funding (e.g., Wellcome Trust). This is emblematic of broader international efforts to measure and reduce carbon emissions [18]. For example, the Carbon Disclosure Project [19], [20], [21] and ISO standards on climate neutrality [22] exemplify the rising global pressure for emissions disclosure across all levels of society. Sustainability is thus becoming an urgent, unavoidable priority for researchers with many institutions adopting policies which incentivise emissions disclosure and reporting.

The rising demand for sustainability-conscious research poses a series of challenges, predominately the need for usable and trustworthy emissions reporting tools as part of a wider goal of evaluating sustainability of research activities. The rapid drive towards improving higher education sustainability may falter due to a lack of inclusive organisational structures[23], [24], concerns of credibility [25], and an absence of easy-to-use, time and resource inexpensive sustainability tools [26]. A parallel can be drawn between the early rise of corporate sustainability-related reporting in 2006-9 [19]. The adoption, standardisation, and refinement of environment, social, and governance (ESG) policies precipitated large-scale improvement to corporate sustainability culture, optimism on the social benefits of their activities, and improved economic and social output performance. While there is a rise of sustainability action in higher educational institutions, there is a serious and consistent risk of performative displays and umbrella commitments [27]. The outsourcing, top-down control of sustainability auditing separates researchers from the direct consequences of their work. Consequently, we risk dampening the direct, positive improvements to sustainability created by individual adjustment to behaviour [28]. Resources that accurately and directly reinforce to researchers the sustainability consequences of their work, and can evidence researcher’s achievements in improving their sustainability, are vital to fostering and sustaining a culture of transparent sustainability awareness and active sustainability improvement.

A growing landscape of sustainability tools offers a rich opportunity for research groups to be pioneers within their fields by transparently reporting their research’s sustainability within publications. Accreditation programs can offer integrated sustainability calculators (e.g., LEAF) which provides an optional, route focused on emissions savings (e.g., saving of emissions from changing waste processes)[3]. Several carbon footprinting tools for computation have been developed (i.e., Green Algorithms [29], CodeCarbon [30], CarbonTracker [31]) which offer hardware contextualised energy-to-emission estimation targeted at computation-heavy research. However, such calculators, while useful, rely on generic emissions conversion, largely ignoring the region- and time-specific factors that affect electricity grids which, in turn, modulate conversion factors [32]. Further, tools like GES 1point5 [33] offer carbon accounting at the level of a research institution for group’s travel and laboratory-wide electricity usage but are exclusively based on French emissions factors, and do not cater to individual, lab-specific activities. While current carbon footprinting tools offer powerful institutional-wide, aggregated carbon footprinting, there are few tools providing real-time, session-based assessment of lab activities that utilise region-specific, time-resolved energy-to-emission (kWh/gCO_2_e) conversation factors. Further, the absence of a single-use platform for comprehensive auditing inhibits uptake and usage of sustainability tools and reporting more generally.

Here, we present WillCO_2_st, a software program that enables research groups across the world to estimate the carbon dioxide emissions equivalent (CO_2_e) for experiments in a given research project. Further, WillCO_2_st enables lab leaders to examine carbon costs across projects within the group, visualising the sustainability across projects and time, and automatically generating carbon emissions focused sustainability statements and reports for papers, grants, and internal audits. To show compatibility with other calculators, we have integrated CodeCarbon [30] into WillCO_2_st to enable computational auditing to sit alongside experimental emissions auditing. In addition, WillCO_2_st offers persistent signalling to experimenters to nudge toward suitable low-emission alternative times to conduct high-energy research. To exemplify the utility of WillCO_2_st, we report on the sustainability of our previous open access publication. Further, we quantify and explore the comparative consequence of shifting the time-of-day that experiments were conducted to lower-energy times, guided by WillCO_2_st’s persistent functionality.

## Results

### Functionality of WillCO_2_st

WillCO_2_st is a free software tool that enables researchers to produce a sustainability audit for experimental work and offers on-demand planning for low-carbon experimentation (Figure 1). WillCO_2_st is hosted on a single main-lab computer with a PC operating system where the principal investigator, or designated sustainability lead, can create a central lab account and add lab members with personalised logins (Figure 2). WillCO_2_st offers two functions: day-by-day nudging for experimenters to conduct high-energy research at lower carbon times and creation of either prospective or retrospective emissions estimations for projects, grants, or audits. WillCO_2_st makes use of APIs that track carbon intensity of electrical grid usage for given locations: Carbon Intensity API for the United Kingdom [34], [35], Electricity Maps for International locations [32], [36], and RTS éCO2mix for France [37]. These APIs reflect that the fuel mixtures driving electricity production of national grids are not homogenous across geography or time[35], [38]. For instance, the Scottish electricity grid mix includes considerably more renewable energy (i.e., wind, biomass) and non-carbon sources (i.e., nuclear) than the English grid, which still relies more on non-renewable sources of energy (i.e., gas). Further, the demands upon local energy grids fluctuate predictably over time with demand spikes aligning with the end of the work day or holidays depleting renewable sources and battery residuals induces more utilisation of domestic or imported non-renewable-driven electricity [32], [39]. The precision and accuracy of electricity-based emissions reporting (i.e., Scope 2) depend upon having real-time, region-specific energy-to-emissions data. WillCO_2_st integrates free-to-request API databases to assure specificity in scope 2 calculation for all users. Further, we designed WillCO_2_st to alert experimenters when their local electricity grid is relying on higher carbon energy production and display a 48-hour, machine learning-driven projection for their local grid’s future, offering an email-based, nudging reminder tool for low-carbon experiment planning (Figure 2).

**Figure 1.**
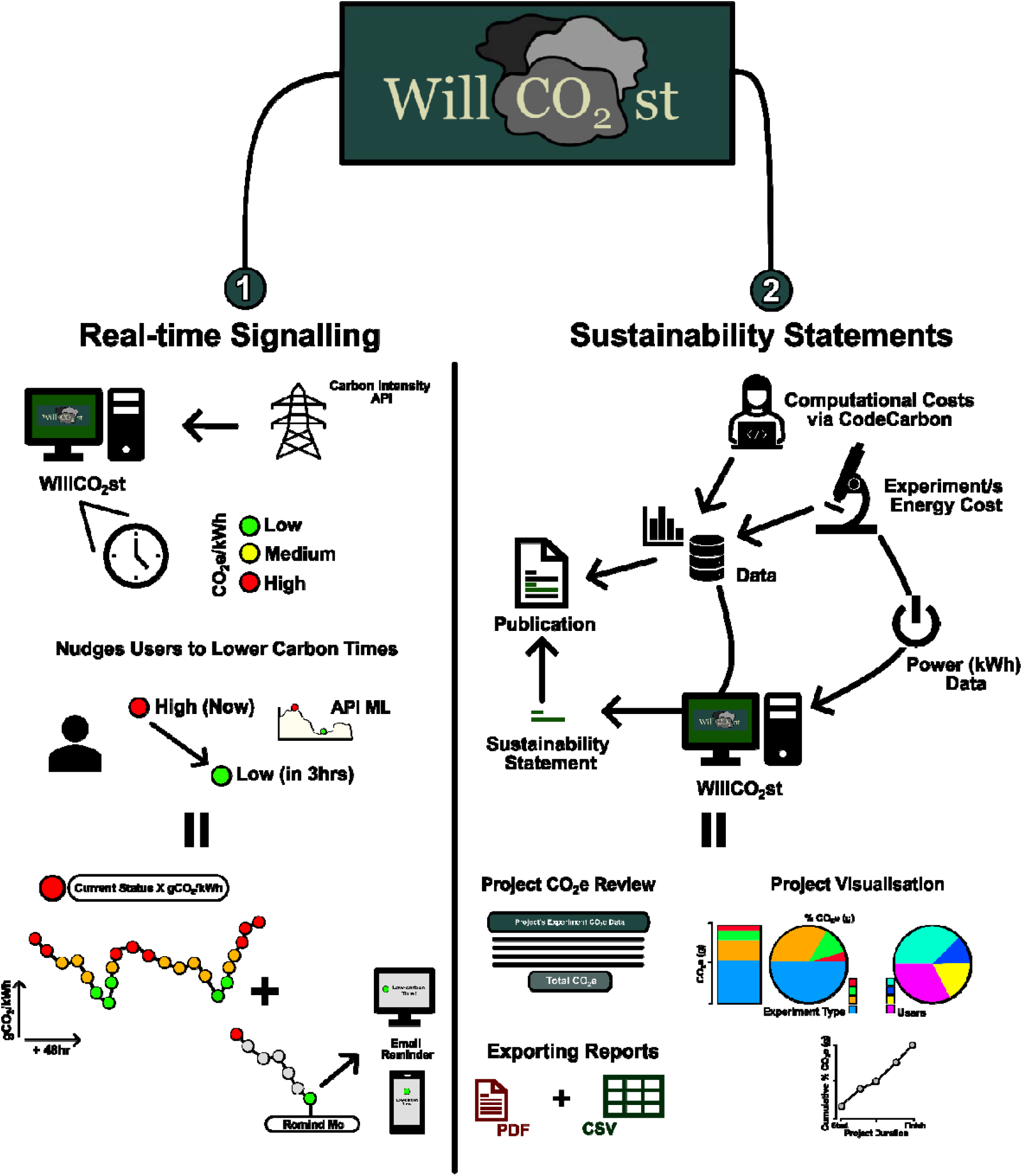
Main Functionality of WillCO_2_st. *WillCO*_2_st’s two main output functions are live grid reporting, enabling future planning for low carbon-time experimental work and the creation of sustainability statements. WillCO_2_st directly reports the local region’s energy (kWh-to-emissions conversion (CO_2_e) factor from an appropriate API the live and 48-hour predicted kWh-to-CO_2_e conversion factor for the research group’s local UK grid network. Users can select lower-carbon times and WillCO_2_st will send an email to remind users to shift activity into low-carbon times to reduce scope 2 emissions. Alongside, WillCO_2_st enables users to create sustainability statements based on user-reported energy consumption levels for experimental equipment. Sustainability statements are generated by WillCO_2_st through the time-of-recording and duration of recording experimental metadata and user-defined power data for each type of experiment (e.g., optogenetics on rig A). WillCO_2_st has integrated CodeCarbon [30] to enable calculations of computational work associated with the project (i.e., Python data extraction, processing, analysis, and visualisation). These sustainability statements are standard text and citation generated by WillCO_2_st to be copy-and-pasted into publications, grants, or reports. WillCO_2_st creates sustainability reports where all users can record and review their project-specific total CO_2_e, visualise the CO_2_e per experiment, user, and cumulative total over the lifetime of the project, and export report for copy-and-pasting sustainability statements into papers (via PDF) and data for open-access reporting (via CSV).

**Figure 2.**
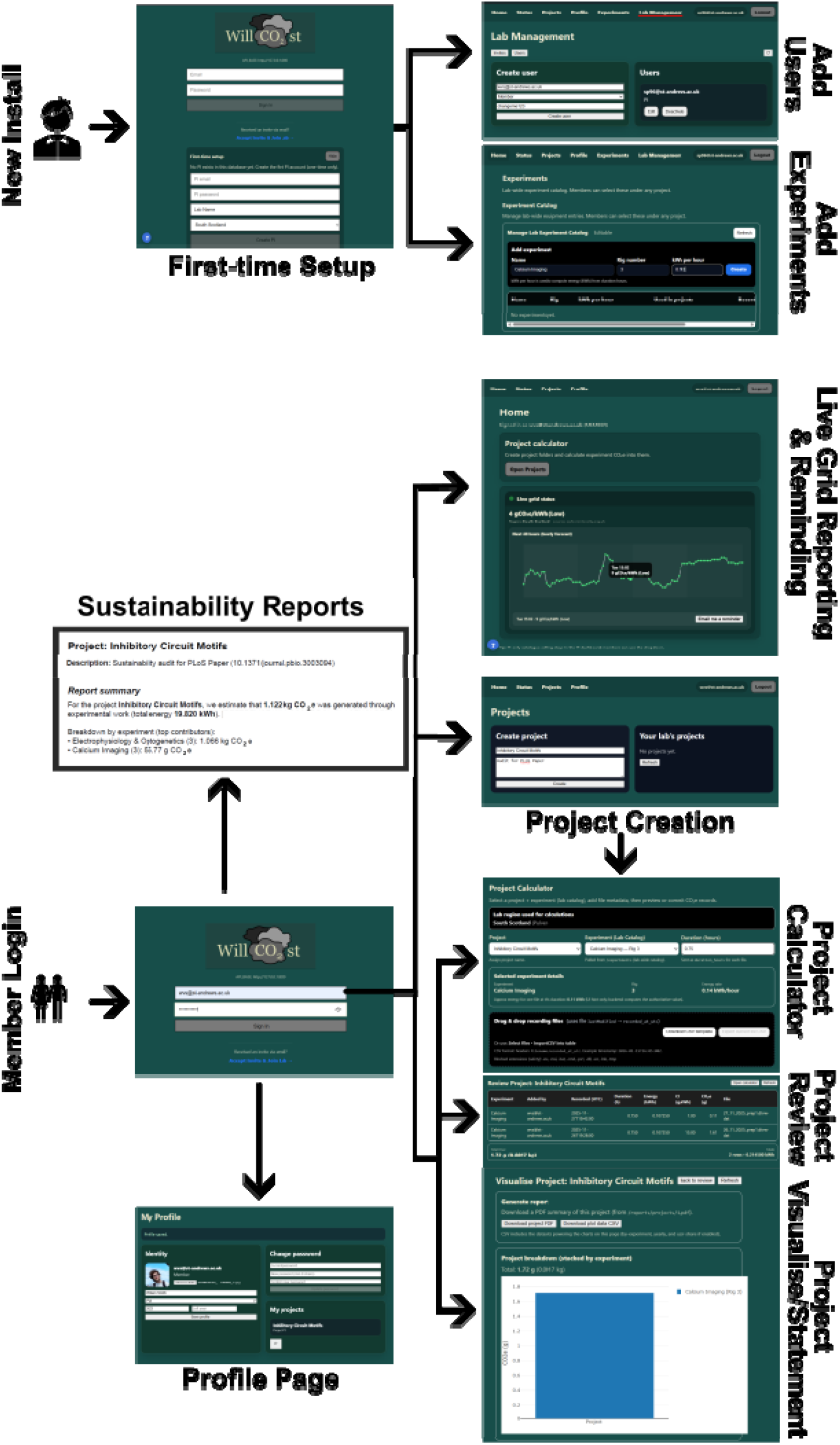
Screenshots of WillCO_2_st Features. WillCO_2_st necessitates local installation and a designated leader (i.e., Lab Head or Sustainability Lead) to create a locally-run account, inviting members to join and populating with experimental energy rating. Members of the WillCO_2_st research group are nudged with live electricity grid status information from their local region and presented with a 48hr projection with an opportunity to email a reminder to the member at low-carbon times. WillCO_2_st facilitates the creation of projects and sustainability audits through calculation, review, visualisation, and reporting pages. Users of WillCO_2_st can edit their own profile offering leaders an ability to record, track, and manage sustainability contributions in the research group.

Alongside real-time signalling, WillCO_2_st can construct sustainability statements for internal audits, methods sections for research articles, grant summaries, and evidence-based drives for improving laboratory sustainability (Figure 1). The rising demand by funding bodies and institutions for sustainability awareness and emissions reduction necessitates accurate, cost- and time-effective methods for internal auditing and reporting. With this in mind, WillCO_2_st enables users to retrospectively audit their experimental work to generate a sustainability statement detailing their experimental carbon footprint. Using real-time energy use recordings, or integrating information from persistent smart meters connected to experimental equipment [35], users can use WillCO_2_st to estimate the generated CO_2_e from their experimental work. WillCO_2_st calculates the CO_2_e for all experimental work in a project by taking the metadata associated with recorded research files and automatically retrospectively searching the historical conversion factors from their local electricity grid as stored in the associated API. Furthermore, we have integrated CodeCarbon [30] – a tool that can compute the CO_2_e associated with computational energy consumption – to provide a more comprehensive experimental emissions audit. With this information, WillCO_2_st can create be used to create carbon-focused sustainability statements that can be exported and added to publications, grant summaries, or internal audits alongside visualisation (e.g., yield and relative experimental contributions) and data compiling for future exploration (Figure 1).

### Using WillCO_2_st

For sustainability audits, WillCO_2_st requires users to specify location, duration, and energy demands of their research to calculate a sustainability statement. Specifically, for experimental work, users declare their research location while setting up their accounts, input the hourly energy consumption (kWh) of their experiments [35], declare the duration of their experiments when updating their projects, and click-and-drag all their primary data files of the work (Figure 3). Experimental energy readings are a crucial component to securing an accurate sustainability estimation for the project’s activities. Given many experimental protocols are heterogenous across research groups (e.g., those involving custom-built microscopes), we advise research groups calculate their own equipment power readings rather than relying on generic reported power values when possible. Here, researchers could use power meters to measure electricity consumption over several hours, days, and experimental repeats to acquire a robust average power reading for each experimental apparatus [35]. Beyond, we would also recommend users invest in dynamic-reporting power meters that report the evolving usage of power from their experimental equipment over time. For computational work, users can enter their Python data files into WillCO_2_st which relies upon CodeCarbon [30] to extract power usage during computation to create a CO_2_e estimate. WillCO_2_st takes the power readings from CodeCarbon, applies the conversion factor of the user’s locate grid, and stores a CO_2_e estimate. A walkthrough of using WillCO_2_st is available in the supplementary information of the manuscript (Supplementary Information Video). The sustainability statement generated using WillCO_2_st can be easily copy-and-pasted into papers under a ‘sustainability statement’ within the method section, or external and internal audits (see Supplementary Information for exemplar Sustainability Statement).

**Figure 3.**
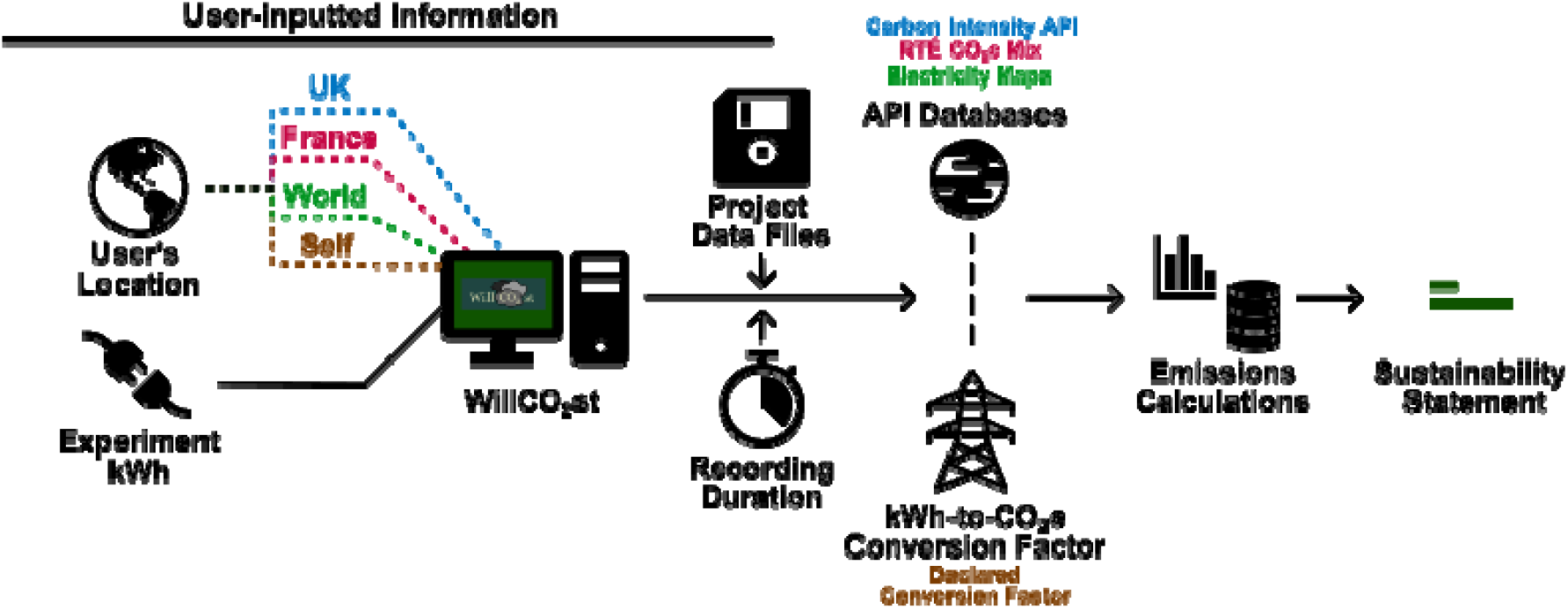
User-inputted Requirements for WillCO_2_st. WillCO_2_st necessitates users specify their research’s location from the options of the UK, France, the World, or, if generated by in-house energy generation, a user-inputted kWh-to-CO_2_e conversion factor. Further, the user must catalogue the power usage of their core experimental equipment which is stored in WillCO_2_st’ experimental catalogue for usage in auditing projects. Users must provide WillCO_2_st will their original raw project data files and specify the duration of the recording in hours. After all pieces of information are available to WillCO_2_st, WillCO_2_st will communicate with the relevant API database for the user’s location to retrospectively calculated the scope 2 emissions for the user’s project and generate a sustainability statement.

In a real-world context, we envisage WillCO_2_st being used for both day-to-day behavioural nudging to conduct experiments at lower carbon times as well as creating sustainability statements before the submission of papers, thesis, grants, and internal audits. Specifically, for sustainability statements, we would recommend using WillCO_2_st to create a new project and uploading the original recording files for each experiment type to create an audit. Indeed, such files form the basis of open-access research and thus should be readily accessible and available to researchers. WillCO_2_st computes the CO_2_e of the experimental research as well as a statement that can be copy-and-pasted into papers, thesis, grants, and internal audits.

#### Sustainability Auditing of Previous Research

To highlight the utility of WillCO_2_st for auditing experimental work, we created a sustainability statement for the experimental work from our most recent publication from the Pulver Lab (Figure 4). In our recent paper [40], we utilised calcium imaging and electrophysiology-optogenetics experiments (N=78) to interrogate the validity of inhibitory modelling approaches to understand rhythmic generation in *Drosophila* (Table 1 in [35]). Briefly, in our previous paper, we imaged the neuronal activity in *Drosophila* ventral nerve cords for 45 minutes in one set of experiments (0.143 kWh) and in another set of experiments manipulated the activity in a restricted set of neurons over 1 hour recordings using electrophysiology and optogenetics techniques (0.426 kWh). Through using those energy readings and by importing all the original recording data files from that project into WillCO_2_st, we estimated that the paper’s experimental work generated 1.48kg CO_2_e (1.42kg CO_2_e for electrophysiology & optogenetics, 0.06kg CO_2_e for calcium imaging) (Figure 4A, left). Given our data (approximately 100GB) was already appropriately organised in line with open-access requirements, the auditing process from importing to WillCO_2_st calculation took < 1 minute. WillCO_2_st calculations for the project were completed in < 1 minute. Expectantly, 94.6% of the experimental footprint occurred pre-submission with reviewer-driven further experiments contributing 80g CO_2_e (5.4%) (Figure 4A, right). While the power usage from our equipment remained stable throughout the recording period (i.e., < ± .1 kWh), if the experimental energy were to fluctuate 10% in each recording period, this would have amounted to ±81.67g CO_2_e variation.

**Figure 4.**
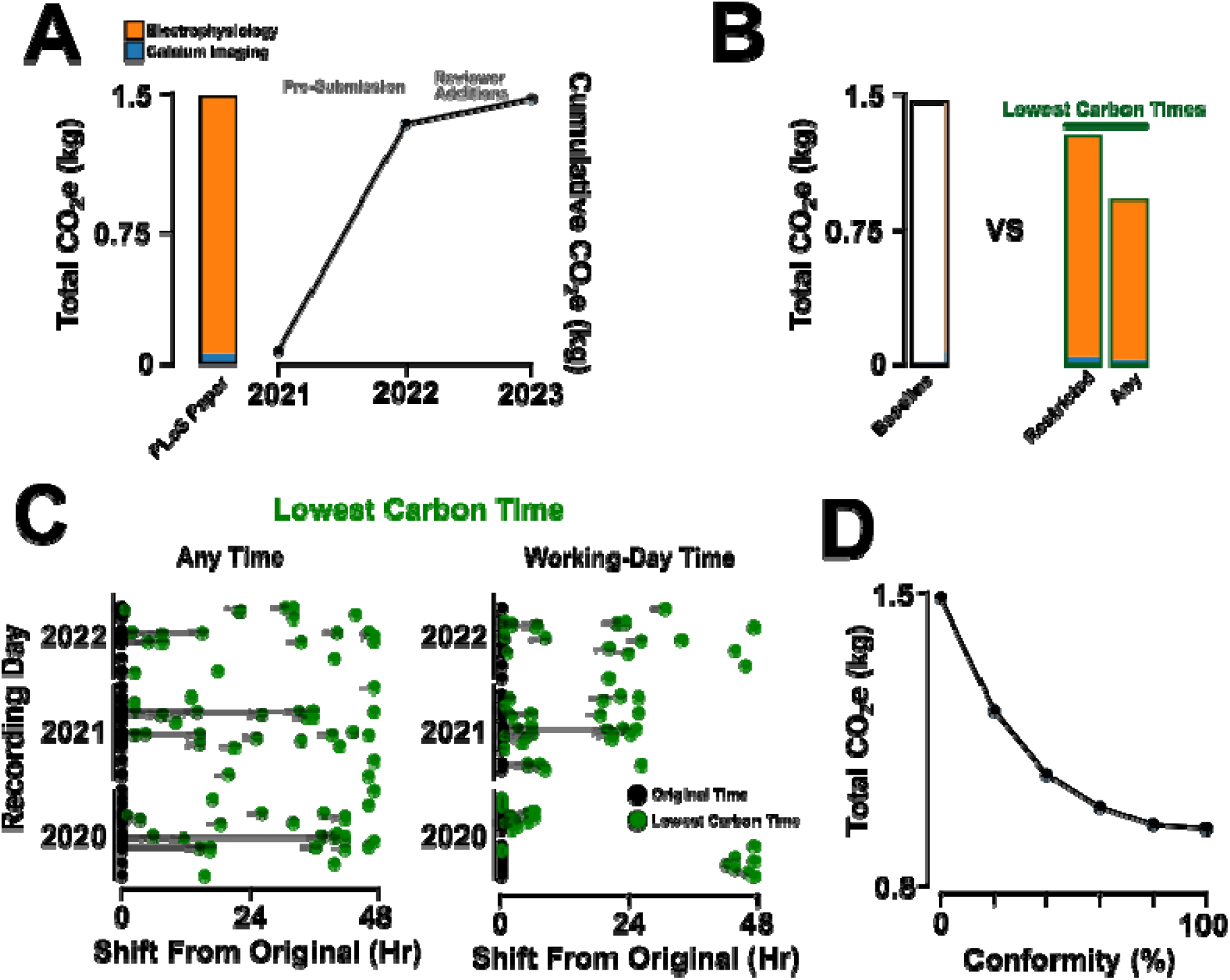
Sustainability Audit for Prior Paper & Consequence of Shifting Conducting Experiments to Reasonable Lowest Carbon Time. (**A**) Carbon footprinting audit by WillCO_2_st for previously published work showing total CO_2_e (kg) and cumulative CO_2_e (kg) between pre-submission (2020-1) to final acceptance after reviewer additions (2023). (**B**) Comparison of total CO_2_e (kg) between actual recorded carbon footprint of experiment work (baseline) to lower carbon times that abide by restricted working hour norms or to any time 48 hours beyond original recording time. (**C**) Scatter shift plot showing the range of lowest carbon times in the South Scotland grid WillCO_2_st, using the Carbon Intensity API, would have recommended the user to conduct the experiment (green dot) compared to original recorded time (black dot) for any time (left) or time within working-day norms (right). (**D**) The effect on reducing the total CO_2_e (kg) generated from the experimental work in the publication based on the % of instances the user shifted to the lowest-carbon time. Note, minimal conformity yields the highest rate of reduction in total CO_2_e (kg) yield from electricity-dependent experimental work.

**Table 1.** Model and energy costs for experimental equipment.

| Type | Equipment | kWh | Model |
| --- | --- | --- | --- |
| Calcium Imaging | Camera and manipulator | 0.045 | Andor iXon3 Camera (100 ms) and MPC-200, TH4-200 Olympus, and ROE-200_1849 Shutter |
|  | Power supply | 0.014 | CAIRN OptoLED Power Supply 3666 |
|  | Perfusion System | 0.002 | Kamoer KCP3-BIOW |
|  | Computer and interface | 0.082 | i7-8700 3.2 GHz CPU, 16 GB RAM, NVIDIA GeForce GTX 1080, and Interface NI-USB-6229 National Instruments Digital I/O System, AD PowerLab Instrument 16/35-PL3616-0154 |
| Electrophysiology and optogenetics | Camera and manipulator | 0.045 | Andor iXon3 Camera and MPC-200, TH4-200 Olympus Microscope Accessory + ROE-200_1849 Shutter Instrument Manipulator |
|  | Power supply | 0.014 | CAIRN OptoLED Power Supply 3666 |
|  | Computer and interface | 0.360 | i7-8700 3.2 GHz CPU, 16 GB RAM, NVIDIA GeForce GTX 1080, and NI-USB-6229 National Instruments Digital I/O System, AD PowerLab Instrument 16/35-PL3616-0154 |
|  | Amplifier | 0.007 | AM System Differential AC Amplifier Model (#59721, 1700) |

**Table 2.**
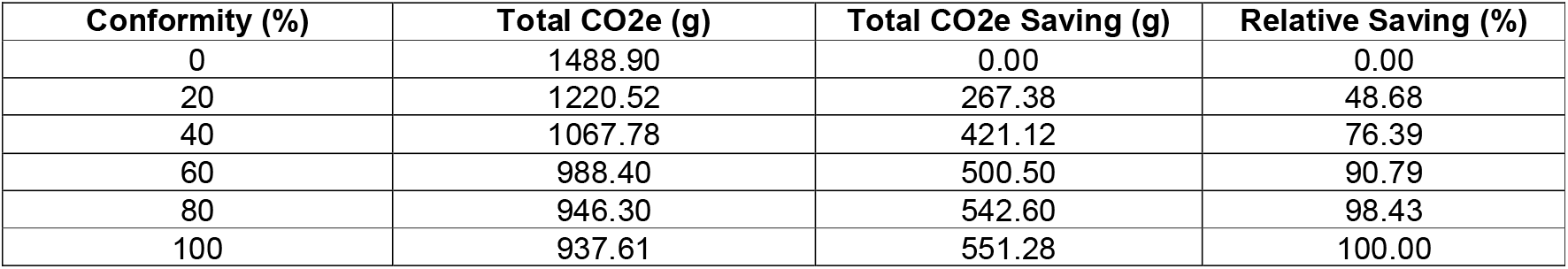
Conformity vs CO_2_e Savings.

To explore the potential utility of WillCO_2_st’s live reporting of conversion data, we assessed the relative effect of changing the time experiments were conducted in our paper to an optimal, lower conversion time on total experimentally-caused carbon load (Figure 4B). To model the effect of rescheduling experimental time, we retrospectively identified the lowest energy-to-CO_2_e conversion time in the South Scotland grid 48 hours post when each experiment for our work was conducted and calculated the relative hypothetical reduction in carbon footprint if we had shifted experiment time. For workhour-constrained rescheduling (Monday-Friday, 9:00AM-20:00PM), we would have witnessed a 12.8% reduction in experiment-derived carbon footprint (1.3kg CO_2_e). If we conformed to workhour-constrained rescheduling recommendations by WillCO_2_st, we would have shifted the start-time of 88.9% of experiments, incurring a 14.66hr (±1.86hr) delay, with 43.1% of experiments now being conducted on a different day than originally performed. We would have most frequently started at either 12:30, 14:00, or 19:00 (24.9% of the time) (Figure 4C, left). In contrast, if we instead conformed to any best-time rescheduling recommendations by WillCO_2_st, we would have shifted the start-time of 100% of experiments, incurring a 26.89hr (±1.85hr) delay, with 80.6% of experiments now being conducted on a different day than originally performed. We would have most frequently started at either 10:00, 12:30, or 02:00 (16.8% of the time) (Figure 4C, right). A 50% relative reduction in total experimental carbon footprint was achievable by accepting and adhering to 21% of WillCO_2_st’s nudges to switch to lower carbon conversion times (Figure 4D, Table 1).

## Discussion

WillCO_2_st is a free Windows-based software tool that empowers any research group to promote conducting experiments at low-carbon times and create, review, and publish sustainability statements for their research papers. Here, we show how WillCO_2_st can effectively and quickly create accurate retrospective sustainability statements for project’s experimental and, through using CodeCarbon [30], computational work. We further show how even partial conformity of shifting experiment time can induce considerable reductions in the carbon footprint of specific experiments. This approach is generalizable and usable across research disciplines. The overall aim with WillCO_2_st is to not just coarsely measure energy usage, but also actively alter how sustainability is implemented and reported at the research group level.

### Promoting research sustainability with WillCO_2_st

The increasing expectations placed upon research institutions and group leaders to be climate conscious and proactive runs a risk of overstretching limited resources and time creating further reporting fatigue [41]. Corporate sustainability reporting (i.e., environmental, social, and governance (ESG) goals) offers a template and insights into the utility, merits, and failings of proactive accountability for carbon-conscious institutions [42]. Fundamentally, current frameworks for sustainability auditing often lack suitable comprehension, sufficient comprehensiveness, widespread inclusion from able stakeholders, specificity to assure compliance and accountable, and viable transparency for both believability and realistic target setting [25], [42], [43], [44], [45], [46], [47].

This work is designed to complement and improve on existing sustainability calculators used by sustainability accreditation schemes such as LEAF. As a gold-accredited LEAF research group, we saw a need for increased accuracy, ease-of-use, and transparency in commonly used laboratory sustainability calculators and designed WillCO_2_st to begin to address these needs. Our goal was to advance the frontier of research sustainability by creating a low-time and energy mechanism for other research groups to self-audit and report on sustainability. We envisage the future landscape of research sustainability incorporating individual-led auditing and scrutinisation especially given the large variability in research techniques, expectations, and evolutions which make top-down, which can alienate researchers [48], [49].

WillCO_2_st is analogous to a home smart metre but with greater specificity, accuracy, accountability, and reportability. Smart metres are widely used in homes to report real-time energy usage and related costs known to induce more conservative use of energy and improved energy literacy[50]. WillCO_2_st resembles the concept of a smart metre applied to research but extends the utility of the concept beyond current tools in a few key dimensions. Firstly, WillCO_2_st enables specific accounting for projects and individual systems rather than from a lab space. Such specificity is quite crucial to reduce the likelihood of double accounting, necessary for carbon literacy, and adds credibility to endeavours to improve sustainability by connecting researchers with the direct carbon consequences of their work. Secondly, WillCO_2_st integrates live-reported conversion data that is highly specific to the geography and time of the user with full transparency and without financial motivation. Thirdly, WillCO_2_st goes beyond just recording and computing carbon footprint data, by enabling creation of sustainability statements in line with the rising expectation of researchers from internal audits (e.g., LEAF) and funding bodies.

The reliability of the emissions reports depends on the validity of the user-inputted energy readings for their experimental equipment. While we found low variability in energy readings for our equipment, acquiring continuous reporting of energy values by power metres is essential to ensure the reliability of WillCO_2_st emissions reports. Thus, careful consideration needs to be paid to all elements necessary for a researcher’s experimental set-up to function (e.g., cooling system, power set-up, conversion instruments). Further, while high-energy pieces of equipment or facilities preclude simple power-metre assessments as described here, such equipment will have power control information available through consultation with technical and estate staff at the research institution. Sustainability statements and carbon appendices should report the specific experimental equipment and their power ratings to bolster reliability of emissions reports [35]. Overall, WillCO_2_st’s unique advantage over other tools is reducing the time burden required to create sustainability audits, particularly in the absence of any institutional or discipline-related sustainability information or support.

### WillCO_2_st & Complementary Tools

WillCO_2_st is designed to complement the variety of existing sustainability auditing tools by providing expansions of and credibility to lab practice certification while adding experimental energy use auditing to current lab footprinting tools. An increasing number of academic institutions are subscribing to sustainability auditing frameworks like LEAF and MyGreenLabs [3]. WillCO_2_st offers a way to increase the accuracy and transparency of internal audits (i.e., LEAF) alongside persistent behaviour nudges that reinforce sustainability training (e.g., MyGreenLabs). Auditing tools like GES 1point5 [33] University of Glasgow’s carbon footprint tool [51], and CO2UNV [52] currently offer research groups a mechanism for research carbon footprinting. Unlike these current tools, WillCO_2_st offers a simple, free and accessible route for sustainability auditing at the individual project level by researchers themselves. Further, WillCO_2_st takes advantage of the free academic access to historical and live emissions databases to generate time- and region-specific estimations rather than relying on static, generalised conversion factors. Beyond institutional and experimental sustainability auditing, open-source tools like Green Algorithms [29], CodeCarbon [30], Cloud Carbon Footprint [53], and EcoLogits [54] uniquely offer assessment of computational, AI, and coding-related experimental research. While powerful, these tools alone do not evaluate specific experimental system energy use (e.g. electrophysiology setups, live imaging systems), use region- and time-specific energy-to-emission conversion factors, nudge users into lower carbon times, nor offer an integrated single-platform reporting tool. In effect, WillCO_2_st offers researchers a complementary and synergistic tool for specialised auditing methodologies that is tailored to unique research disciplines and demands. The validity of any carbon emission is only as valid as the underlying energy and emission factor data. WillCO_2_st depends upon two data sources – user-reporting energy levels with error margins and external-reported emissions factors – whose accuracy remains out of WillCO_2_st’s direct control. However, the risks of these dependencies are mitigated by the transparent APIs we chose to integrate within WillCO_2_st; our sensitivity analysis also shows that the potential for drastic miscalculation is relatively low [30], [32], [34], [36], [37], Obviously, the effectiveness of any tool relies on the user. Researchers must do their own due diligence and carefully consider the utility, viability, applicability and reliability of any tool they use for sustainability reporting.

### Future of Research Sustainability

Transitioning to more sustainable working is a global trend. Researchers are not exempt and they will most likely be progressively required to demonstrate commitment to sustainability in their research practices Pioneering research groups have already begun to offer evidence-based changes to discipline-specific practices in response to calls to promote sustainability [29], [33], [35], [55], [56], [57]. WillCO_2_st provides one tool amongst several valuable resources to promote individual sustainability auditing. For the near future, all experimental and research work will necessitate some level of non-renewable resources. The process of auditing maximises the efficiency and utility of the researcher’s resources while offering direct evidence of engaging with sustainability, as increasingly demanded by institutions and funding bodies. As we show, even partial conformity to lower carbon intensity times can have notable reductions in non-renewable electricity usage and thus can provide unambiguous evidence of sustainability engagement. Importantly, our research in Scotland is performed using a renewable energy grid, as a result, the raw saved CO_2_e value reported here may seem inconsiderable. However, the level of savings would increase in more non-renewable grids [35]. Cumulatively, the potential for large scale energy savings with minimal behavioural change are high.

The future of research sustainability is bright with several, synergistic tools and verifiable methodologies for sustainability accounting. However, the strength of any emissions reporting tool depends entirely on the continued free, open-source access to emissions and conversion data. WillCO_2_st depends upon free access to the data provided by the Carbon Intensity API [34], Electricity Maps [32], [36], and RTS éCO_2_mix [37]. The high uptake and ultimate success of emission-reporting tools depend on the continued free access to credible sources of emission conversion information, requiring developers to be transparent about data collection, analysis, and reporting. Specifically, resources like the ENTSO-E Transparency Platform’s energy generation mix data (https://transparency.entsoe.eu/) form a valid, transparent basis of future, free, accurate emissions models providing deeper EU-wide integration with WillCO_2_st. More broadly, to assure persistent positive improvement in research sustainability, decentralised and democratic methods that empower all stakeholders to monitor, publish and improve their performance is essential to endanger collective action and scrutiny.

Research sustainability will most likely be predominately driven by top-down reporting and auditing requirements as evidenced by the rise of LEAF, MyGreenLabs [3] and the evolving UKRI-led SPARKShub (https://sparkhub.eu/). Thus, the ease-of-use and accuracy of tools that researchers rely upon when completing audits and responding to sustainability demands is essential to drive positive and meaningful improvements in sustainability. A significant impediment to sustainability uptakes remains reporting fatigue, particularly in the face of an ever-increasing workload placed onto academics and researchers [46]. With this in mind, WillCO_2_st is built to streamline emissions reporting and offers a transparent mechanism to signal sustainability engagement. Beyond, WillCO_2_st reduces the fatigue in reporting by consolidating a significant area of research in a stepwise fashion. While not an exhaustive tool, WillCO_2_st is designed to allow researchers to clearly demonstrate engagement with sustainability. From historical audits, showing lab-wide participation, to tracking the success of behavioural nudges – all of these outcomes align with the highest accreditation levels of LEAF and are the core vision of MyGreenLabs: a sustainability-conscious and sustainability-active researcher.

## Methods

### Structure & Dependencies of WillCO_2_st

WillCO_2_st (v1.0) is built using Python (version 3.14.2) and JavaScript using Microsoft Visual Studio 2022. The version of WillCO_2_st reported here exclusively functions within the Windows operating system. WillCO_2_st requires a persistent internet connection to actively communicate with the relevant emissions databases. All packages and dependencies are directly imported by WillCO_2_st within the installer function.

### Conversion Calculations in WillCO_2_st

WillCO_2_st (v1.0) rests upon historical and active conversion reporting by the Carbon Intensity API for UK researchers [34], Electricity Maps for global researchers [32], [36], and RTS éCO_2_mix for French researchers [37]. These API offers access to historical, live, and prospective energy mixture data for electricity grid to compute dynamic, region-specific, and time-specific kWh-to-CO_2_e conversion factors. WillCO_2_st accesses these databases for live grid reporting, prospective grid reporting, and historical sustainability auditing. Users can input energy readings utilising cheap, accessible power meters [35] into WillCO_2_st that then computes the CO_2_e via time-specific kWh-to-CO_2_e conversion factors reported by the APIs. The time-of-recording is either imputable by the user as a the direct, personally-recorded time of recording (via CSV) or via automatic recognition of ‘last-modified’ metadata in the saved file if the user chooses to click-and-drag (Figure 1’s ‘Add Experiments’, Figure 3, see Supplementary Video and User Guide). The user is cautioned that ‘last-modified’ does not axiomatically mean precise recording time, so users are provided a .CSV option to import recording dates of datafiles if datafiles are modified post recording.

### Algorithm Determining Experiment Change-of-time Variation Impact

Using a custom-build Python script (available in our data repository) and historical data from the emission factor APIs, we determined the variation impact of changing experiment time by identifying the lowest kWh-to-CO_2_e conversion factor within a 48-hour window post the original experiment time. The last modified and original recording time metadata for all experimental files was used to determine the precise 30-minute window of all experiment’s original recording time. We either applied no time restrictions and retrieved the lowest kWh-to-CO_2_e conversion factor (“Any”) or the lowest kWh-to-CO_2_e conversion factor within working hour restrictions (i.e., excluding weekends and operating exclusively between 9:00AM and 20:00PM). We adjusted the counterfactual recording times to prevent impossible overlap in the recording session. The final recorded CO_2_e was calculated as the power rating of the experimental set-up multiplied by the retrieved conversion factor. We assumed the power ratings of our experimental equipment [35] remained consistently across the duration of the project and any degradation would have had a minimal impact on final power usage.

## Supplementary Figure

**Supplementary Figure 1.**
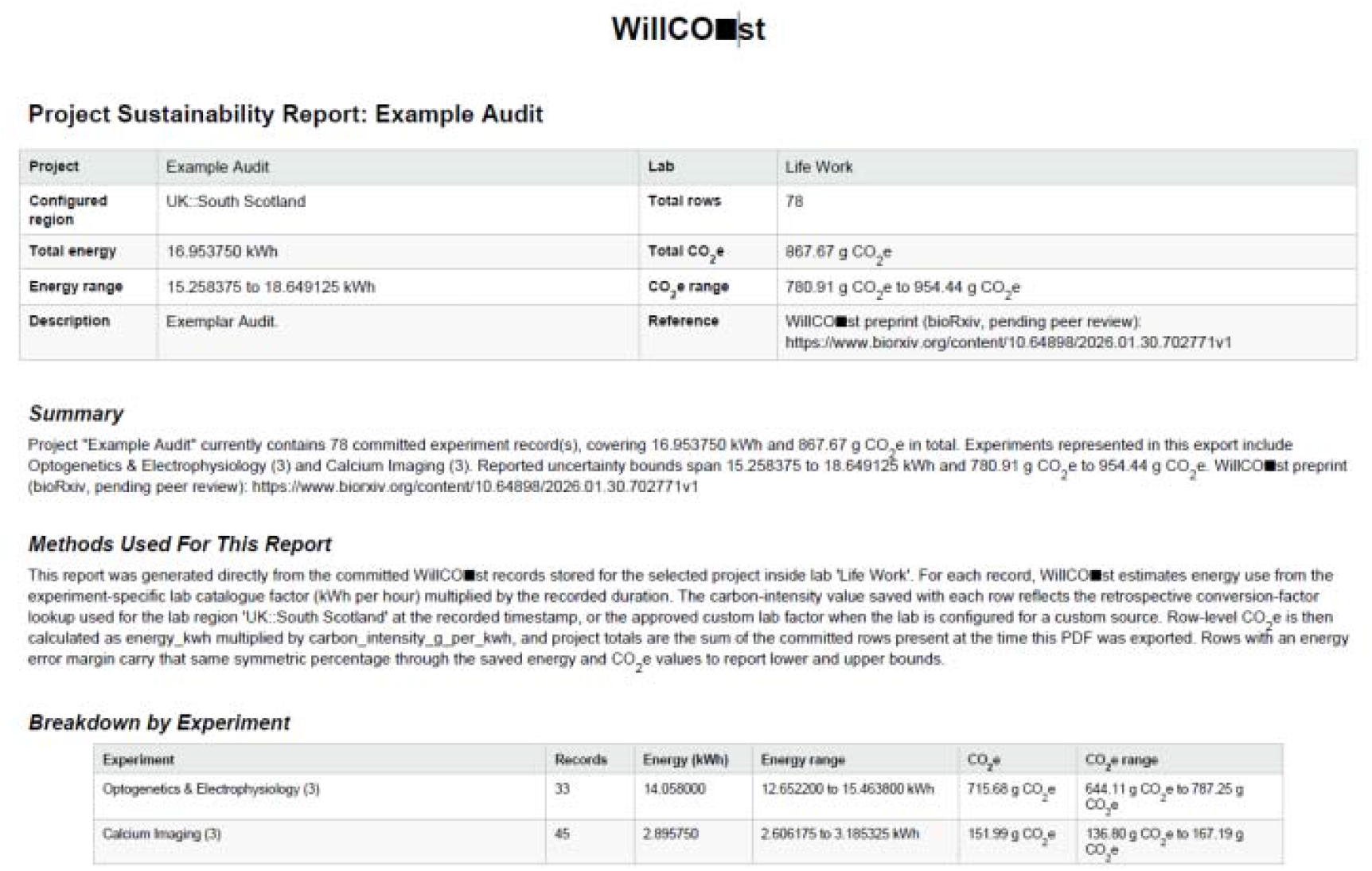
Exemplar Sustainability Audit from WillCO_2_st.

## Acknowledgements

This project was made possible by funding from Scotland’s Future Series (University of St Andrews) to W.V.S.; an Industrial CASE PhD studentship (UKRI Biotechnology and Biological Sciences Research Council [BBSRC] grant number BB/T00875X/1) to W.V.S. The historical data underpinning WillCO_2_st depends on the continued free, universal, open-access to the listed publicly-available APIs.

## Author Contributions

W.V.S. conceived the study. W.V.S. designed, created, and tested WillCO_2_st. W.V.S performed all analysis and drafted the manuscript. W.V.S and S.R.P revised the manuscript. S.R.P. supervised the project.

## Data Availability

The research data underpinning this publication can be accessed at: <u>10.17630/788c0aea-ab66-43e0-a7a5-761b5c64c963</u>.

## References

[1] UKRI, ‘Concordat for the Environmental Sustainability of Research and Innovation Practice’, 2022. doi: https://www.ukri.org/about-us/what-we-do/environmental-sustainability/concordat-for-the-environmental-sustainability-of-research-and-innovation-practice/.

[2] M. De Paepe, L. Jeanneau, J. Mariette, O. Aumont, and A. Estevez-Torres, ‘Purchases dominate the carbon’, PLOS Sustain. Transform., vol. 3, no. 7, p. e0000116, Jul. 2024, doi: 10.1371/journal.pstr.0000116.

[3] B. R. Schell and N. Bruns, ‘Lab sustainability programs LEAF and My Green Lab®: impact, user experience & suitability’, RSC Sustain., vol. 2, no. 11, pp. 3383–3396, 2024, doi: 10.1039/D4SU00387J.

[4] Wellcome Trust, ‘Wellcome Trust Environmental Sustainability’, 2025. [Online]. Available: https://wellcome.org/research-funding/guidance/policies-grant-conditions/environmental-sustainability-funding-policy

[5] CRUK, ‘Environmental sustainability in research’, 2025. [Online]. Available: https://www.cancerresearchuk.org/funding-for-researchers/how-we-deliver-research/environmental-sustainability-in-research

[6] NIHR, ‘Sustainability and net zero’, 2025. [Online]. Available: https://www.nihr.ac.uk/about-us/our-contribution-to-research/sustainability-and-net-zero.htmL

[7] British Heart Foundation, ‘Environmental sustainability’, 2025. [Online]. Available: https://www.bhf.org.uk/what-we-do/how-we-work/environmental-sustainability

[8] Royal Academy of Engineering, ‘Environmental sustainability strategy’, 2025. [Online]. Available: https://raeng.org.uk/about-us/sustainability

[9] European Commission, ‘Horizon Europe Work Programme 2025’. [Online]. Available: https://ec.europa.eu/info/funding-tenders/opportunities/docs/2021-2027/horizon/wp-call/2025/wp-12-missions_horizon-2025_en.pdf

[10] FDFA/DETEC, ‘2030 Sustainable Development Strategy’. [Online]. Available: https://www.agenda-2030.eda.admin.ch/en/implementing-in-switzerland

[11] Deutsche and Forschungsgemeinschaft, ‘Anchoring sustainability considerations in DFG funding activities’. [Online]. Available: https://www.dfg.de/resource/blob/334032/empfehlungen-en.pdf

[12] R. Guérin, A. McGovern, and K. Nahrstedt, ‘Report on the NSF Workshop on Sustainable Computing for Sustainability (NSF WSCS 2024)’, 2024, arXiv. doi: 10.48550/ARXIV.2407.06119.

[13] National Institute of Environmental Health Sciences, ‘Strategic Plan 2025-2029’. [Online]. Available: https://www.niehs.nih.gov/about/strategicplan

[14] Australian Research Council, ‘Annual Report 2024-25’. [Online]. Available: https://www.arc.gov.au/sites/default/files/2025-10/ARC%20Annual%20Report%202024-25.pdf

[15] Science Foundation Ireland, ‘Climate Action Strategy 2024-2027’. [Online]. Available: https://www.sfi.ie/research-news/publications/SFI-Climate-Strategy.pdf

[16] European Commission, ‘Marie Skłodowska-Curie Actions Green Charter’. [Online]. Available: https://op.europa.eu/en/publication-detail/-/publication/d8e5b75a-ae2d-11f0-89c6-01aa75ed71a1/language-en#

[17] Royal Academy of Chemistry, ‘Environmental sustainability strategy’, 2025. [Online]. Available: https://www.rsc.org/prizes-funding/funding/sustainable-labs-grants/

[18] A. Di Vaio, M. Chhabra, A. Zaffar, and D. Balsalobre-Lorente, ‘Accounting and Accountability in the Transition to Zero-Carbon Energy for Climate Change: A Systematic Literature Review’, Bus. Strategy Environ., vol. 34, no. 5, pp. 5925–5946, Jul. 2025, doi: 10.1002/bse.4282.

[19] Z. Borghei, ‘Carbon disclosure: a systematic literature review’, Account. Finance, vol. 61, no. 4, pp. 5255–5280, Dec. 2021, doi: 10.1111/acfi.12757.

[20] K. Alsaifi, M. Elnahass, and A. Salama, ‘Carbon disclosure and financial performance: UK environmental policy’, Bus. Strategy Environ., vol. 29, no. 2, pp. 711–726, Feb. 2020, doi: 10.1002/bse.2426.

[21] F. Chen, M. Hussain, J. A. Khan, G. M. Mir, and Z. Khan, ‘Voluntary disclosure of greenhouse gas emissions by cities under carbon disclosure project: A sustainable development approach’, Sustain. Dev., vol. 29, no. 4, pp. 719–727, Jul. 2021, doi: 10.1002/sd.2169.

[22] M. Finkbeiner, A. Inaba, R. Tan, K. Christiansen, and H.-J. Klüppel, ‘The New International Standards for Life Cycle Assessment: ISO 14040 and ISO 14044’, Int. J. Life Cycle Assess., vol. 11, no. 2, pp. 80–85, Mar. 2006, doi: 10.1065/lca2006.02.002.

[23] R. Sassen, D. Dienes, and J. Wedemeier, ‘Characteristics of UK higher education institutions that disclose sustainability reports’, Int. J. Sustain. High. Educ., vol. 19, no. 7, pp. 1279–1298, Nov. 2018, doi: 10.1108/IJSHE-03-2018-0042.

[24] R. Sassen and L. Azizi, ‘Assessing sustainability reports of US universities’, Int. J. Sustain. High. Educ., vol. 19, no. 7, pp. 1158–1184, Nov. 2018, doi: 10.1108/IJSHE-06-2016-0114.

[25] M. Pramono Sari, F. Faisal, and P. Harto, ‘The determinants of higher education institutions’ (HEIs) sustainability reporting’, Cogent Bus. Manag., vol. 10, no. 3, p. 2286668, Dec. 2023, doi: 10.1080/23311975.2023.2286668.

[26] J. Abello-Romero, C. Mancilla, K. Restrepo, W. Sáez, I. Durán-Seguel, and F. Ganga-Contreras, ‘Sustainability Reporting in the University Context—A Review and Analysis of the Literature’, Sustainability, vol. 16, no. 24, p. 10888, Dec. 2024, doi: 10.3390/su162410888.

[27] B. Latter and S. Capstick, ‘Climate Emergency: UK Universities’ Declarations and Their Role in Responding to Climate Change’, Front. Sustain., vol. 2, p. 660596, May 2021, doi: 10.3389/frsus.2021.660596.

[28] R. Zhou, ‘How UK universities approach sustainability: A timely review’, J. Adult Contin. Educ., vol. 31, no. 1, pp. 54–73, Mar. 2025, doi: 10.1177/14779714241240985.

[29] L. Lannelongue, J. Grealey, and M. Inouye, ‘Green Algorithms: Quantifying the Carbon Footprint of Computation’, Adv. Sci., vol. 8, no. 12, p. 2100707, Jun. 2021, doi: 10.1002/advs.202100707.

[30] R. Fischer, ‘Ground-Truthing AI Energy Consumption: Validating CodeCarbon Against External Measurements’, 2025, arXiv. doi: 10.48550/ARXIV.2509.22092.

[31] L. F. W. Anthony, B. Kanding, and R. Selvan, ‘Carbontracker: Tracking and Predicting the Carbon Footprint of Training Deep Learning Models’, 2020, arXiv. doi: 10.48550/ARXIV.2007.03051.

[32] G. Boyle, Ed., Renewable electricity and the grid: the challenge of variability, Online-Ausg. London Sterling, VA: Earthscan, 2007.

[33] J. Mariette et al., ‘An open-source tool to assess the carbon footprint of research’, Environ. Res. Infrastruct. Sustain., vol. 2, no. 3, p. 035008, Sep. 2022, doi: 10.1088/2634-4505/ac84a4.

[34] National Energy System Operator (NESO), ‘Carbon Intensity API’. [Online]. Available: https://carbonintensity.org.uk/

[35] W. V. Smith, A. Bebbington, R. Sircar, M. C. Gather, and S. R. Pulver, ‘Students as Carbon Accountants: Calculating Carbon Costs of a PhD in Neuroscience’, GENETICS, p. iyaf268, Dec. 2025, doi: 10.1093/genetics/iyaf268.

[36] Electricity Maps, ‘Electricity Maps API & methodology’. [Online]. Available: https://www.electricitymaps.com/

[37] RTE (Reseau de Transport d’Electricite), ‘Eco2mix - electricity generation and CO2 emissions data’. [Online]. Available: https://eco2mix.rte-france.com/curves/eco2mixWeb

[38] M. Lockwood, ‘Transforming the grid for a more environmentally and socially sustainable electricity system in Great Britain is a slow and uneven process’, Proc. Natl. Acad. Sci., vol. 120, no. 47, p. e2207825120, Nov. 2023, doi: 10.1073/pnas.2207825120.

[39] M. Asif, J. Currie, and T. Muneer, ‘The role of renewable and non-renewable sources for meeting future UK energy needs’, Int. J. Nucl. Gov. Econ. Ecol., vol. 1, no. 4, p. 372, 2007, doi: 10.1504/IJNGEE.2007.016657.

[40] J. Francis et al., ‘Inhibitory circuit motifs in Drosophila larvae generate motor program diversity and variability’, PLOS Biol., vol. 23, no. 4, p. e3003094, Apr. 2025, doi: 10.1371/journal.pbio.3003094.

[41] M. De Jong, ‘Sustainability reporting: can we avoid another triumph of hope over experience?’, J. Public Budg. Account. Financ. Manag., vol. 37, no. 6, pp. 354–369, Dec. 2025, doi: 10.1108/JPBAFM-01-2025-0024.

[42] S. Ramanathan and R. Isaksson, ‘Sustainability reporting as a 21st century problem statement: using a quality lens to understand and analyse the challenges’, TQM J., vol. 35, no. 5, pp. 1310– 1328, Jun. 2023, doi: 10.1108/TQM-01-2022-0035.

[43] B. O’Dwyer and D. L. Owen, ‘Assurance statement practice in environmental, social and sustainability reporting: a critical evaluation’, Br. Account. Rev., vol. 37, no. 2, pp. 205–229, Jun. 2005, doi: 10.1016/j.bar.2005.01.005.

[44] Y. P. Lopez and W. F. Martin, ‘University Mission Statements and Sustainability Performance’, Bus. Soc. Rev., vol. 123, no. 2, pp. 341–368, Jun. 2018, doi: 10.1111/basr.12144.

[45] A. Wiek, L. Withycombe, and C. L. Redman, ‘Key competencies in sustainability: a reference framework for academic program development’, Sustain. Sci., vol. 6, no. 2, pp. 203–218, Jul. 2011, doi: 10.1007/s11625-011-0132-6.

[46] B. A. Christie, K. K. Miller, R. Cooke, and J. G. White, ‘Environmental sustainability in higher education: What do academics think?’, Environ. Educ. Res., vol. 21, no. 5, pp. 655–686, Jul. 2015, doi: 10.1080/13504622.2013.879697.

[47] L. White and B. F. Noble, ‘Strategic environmental assessment for sustainability: A review of a decade of academic research’, Environ. Impact Assess. Rev., vol. 42, pp. 60–66, Sep. 2013, doi: 10.1016/j.eiar.2012.10.003.

[48] R. Sanchez, J. Galbreath, and G. Nicholson, ‘Building Sustainability Competence from the Top Down: A Model for Researching and Improving Boards of Directors’ Influence on Firms’ Sustainability Performance’, in Mid-Range Management Theory: Competence Perspectives on Modularity and Dynamic Capabilities, R. Sanchez, A. Heene, S. Polat, and U. Asan, Eds, Emerald Publishing Limited, 2017, pp. 69–107. doi: 10.1108/S1744-211720170000008004.

[49] C. Jungwirth and E. Mueller, ‘The Sustainability of Clusters - Consequences of Different Governance Regimes of Top-Down and Bottom-Up Cluster Initiatives’, SSRN Electron. J., 2010, doi: 10.2139/ssrn.1615380.

[50] N. Mogles et al., ‘How smart do smart meters need to be?’, Build. Environ., vol. 125, pp. 439–450, Nov. 2017, doi: 10.1016/j.buildenv.2017.09.008.

[51] Centre for Sustainable Solutions, ‘Glasgow Carbon Footprint Tool for Researchers’, 2025. [Online]. Available: https://www.gla.ac.uk/research/az/sustainablesolutions/ourprojects/carbonfootprinttool/

[52] K. Valls-Val and M. D. Bovea, ‘Carbon footprint assessment tool for universities: CO2UNV’, Sustain. Prod. Consum., vol. 29, pp. 791–804, Jan. 2022, doi: 10.1016/j.spc.2021.11.020.

[53] Thoughtworks, ‘Cloud Carbon Footprint’, 2025. [Online]. Available: https://www.cloudcarbonfootprint.org/docs/

[54] ‘Ecologits Calculator’, 2025. [Online]. Available: https://huggingface.co/spaces/genai-impact/ecologits-calculator

[55] T. Spruell, H. Webb, Z. Steley, J. Chan, and A. Robertson, ‘Environmentally sustainable emergency medicine’, Emerg. Med. J., vol. 38, no. 4, pp. 315–318, Apr. 2021, doi: 10.1136/emermed-2020-210421.

[56] N. E. Souter et al., ‘Measuring and reducing the carbon footprint of FMRI preprocessing in FMRIPREP’, Hum. Brain Mapp., vol. 45, no. 12, p. e70003, Aug. 2024, doi: 10.1002/hbm.70003.

[57] N. E. Souter et al., ‘Ten recommendations for reducing the carbon footprint of research computing in human neuroimaging’, Imaging Neurosci., vol. 1, pp. 1–15, Dec. 2023, doi: 10.1162/imag_a_00043.

